# A web-based platform of nucleotide sequence alignments of plants

**DOI:** 10.1101/617035

**Authors:** Chiara Santos, João Carneiro, Filipe Pereira

**Affiliations:** Interdisciplinary Centre of Marine and Environmental Research (CIIMAR), University of Porto, Terminal de Cruzeiros do Porto de Leixões; Avenida General Norton de Matos, S/N 4450-208 Matosinhos – Portugal; University of Porto, Praça Gomes Teixeira 4099-002 Porto, Portugal

**Author notes:** These authors contributed equally to this work (Chiara Santos, João Carneiro). Corresponding author: F. Pereira.

**Keywords:** DNA sequences, Multiple sequence alignments, Plant families

## Abstract

In recent years, a large number of nucleotide sequences have become available for plant species by the advent of massive parallel sequencing. The use of genomic data has been important for agriculture, food science, medicine or ecology. Despite the increasing amount of data, nucleotide sequences are usually available in public databases as isolated records with some descriptive information. Researchers interested in studying a wide range of specific plant families are forced to do multiple searches, sequence downloads, data curation and sequence alignments. In order to help researches overcoming these problems, we have built a comprehensive on-line resource of curated nucleotide sequence alignments for plant research, named PlantAligDB (available at http://plantaligdb.portugene.com). The latest release incorporates 514 alignments with a total of 66,052 sequences from six important genomic regions: *atp*F-*atp*H, *psb*A-*trn*H, *trn*L, *rbc*L, *mat*K and ITS. The alignments represent 223 plant families from a variety of taxonomic groups. The users can quickly search the database, download and visualize the curated alignments and phylogenetic trees using dynamic browser-based applications. Different measures of genetic diversity are also available for each plant family. We also provide the workflow script that allows the user to do the curation process, explaining the steps involved. Overall, the PlantAligDB provides a complete, quality checked and regularly updated collection of alignments that can be used in taxonomic, DNA barcoding, molecular genetics, phylogenetic and evolutionary studies.

## Introduction

The recent development of high-throughput sequencing technologies has significantly increased the number of nucleotide sequences available in public databases (Egan *et al*. 2012; Feuillet *et al*. 2011). Complete genome sequences are now accessible in public databases (e.g., EnsemblPlants) (https://plants.ensembl.org/index.html) for the analysis and visualisation of genomic data for an ever-growing number of plants, such as *Beta vulgaris, Prunus persica* and *Citrus sinensis*, among many others. Sequences from individual genes or gene regions have also been deposit in public databases as a result of international initiatives. For instance, the DNA barcoding project has released thousands of sequences aiming at species identification and taxonomic classification of plants, mostly from the chloroplast DNA (cpDNA) protein-coding genes *rbc*L and *mat*K [CBOL Plant Working Group’ - (Group *et al*. 2009; Hollingsworth *et al*. 2011)]. The plastid *trn*L (UAA) intron is another good example of a cpDNA region highly represented in sequence databanks (Taberlet *et al*. 2007).

Several web-based databases are available for plant genome sequences, usually dedicated to a single species or a genomic feature [e.g., (Lai *et al*. 2012; Meyer *et al*. 2005; Numa & Itoh 2014; Sakai *et al*. 2013)]. However, most nucleotide sequences are accessible in public databases as isolated records with simple descriptive information (taxonomy, geography, publications, etc.). For instance, the NCBI Entrez Nucleotide database (http://www.ncbi.nlm.nih.gov) and the BOLD - The Barcode of Life Data System (www.barcodinglife.org) (Ratnasingham & Hebert 2007) are useful repositories with descriptive information for sequence or species. The TreeBASE (https://www.treebase.org) is a repository of phylogenetic information with user-submitted phylogenetic trees and the data used to generate them. Nevertheless, researchers interested in studying a large number of plant families are forced to do multiple searches and sequence downloads of genetic data for their investigations. Moreover, the available sequences are not aligned and curated. The multiple sequence alignment step is critical because it determines the accuracy of the subsequence analyses, such as phylogenetic inference, identification of conserved motifs, function prediction, etc. Building accurate sequence alignments involves many steps, including sequence file conversion, run of alignment algorithms in local computers or webservers, selection of best alignment parameters, and manual fine-tuning of the alignment. This process is laborious and requires costly computational resources, which are not always available.

We describe an on-line database (PlantAligDB, available at http://plantaligdb.portugene.com) with a comprehensive, automatically curated and regularly updated collection of alignments from diverse plant families (Figure 1), and respective phylogenetic inference of each alignment. The PlantAligDB is a consistent repository of curated alignments and phylogenetic trees that are generated by the same workflow. For example, the Orchidaceae family is frequently referred to as a critical group, whose species are difficult to identify. Because of their ecological importance, ongoing studies continue to be made to reduce their extinction risk and maintain their diversity (Li *et al*. 2018). The PlantAligDB can help researchers designing accurate methods for plant identification (*mat*K and *rbc*L are used in DNA barcoding projects) whether by identification of conserved motives that enable the design of primers. The PlantAligDB alignments can be used as a reference database for phylogenetic studies, allowing the construction of reference phylogenetic trees of the different regions [e.g., genomic regions *atp*F*-atp*H (Domenech *et al*. 2014), *psb*A*-trn*H (Dong *et al*. 2012) and ITS (Karehed *et al*. 2008)]. Moreover, it provides useful data to understand the genetic diversity of the selected genomic regions.

**Figure 1:**
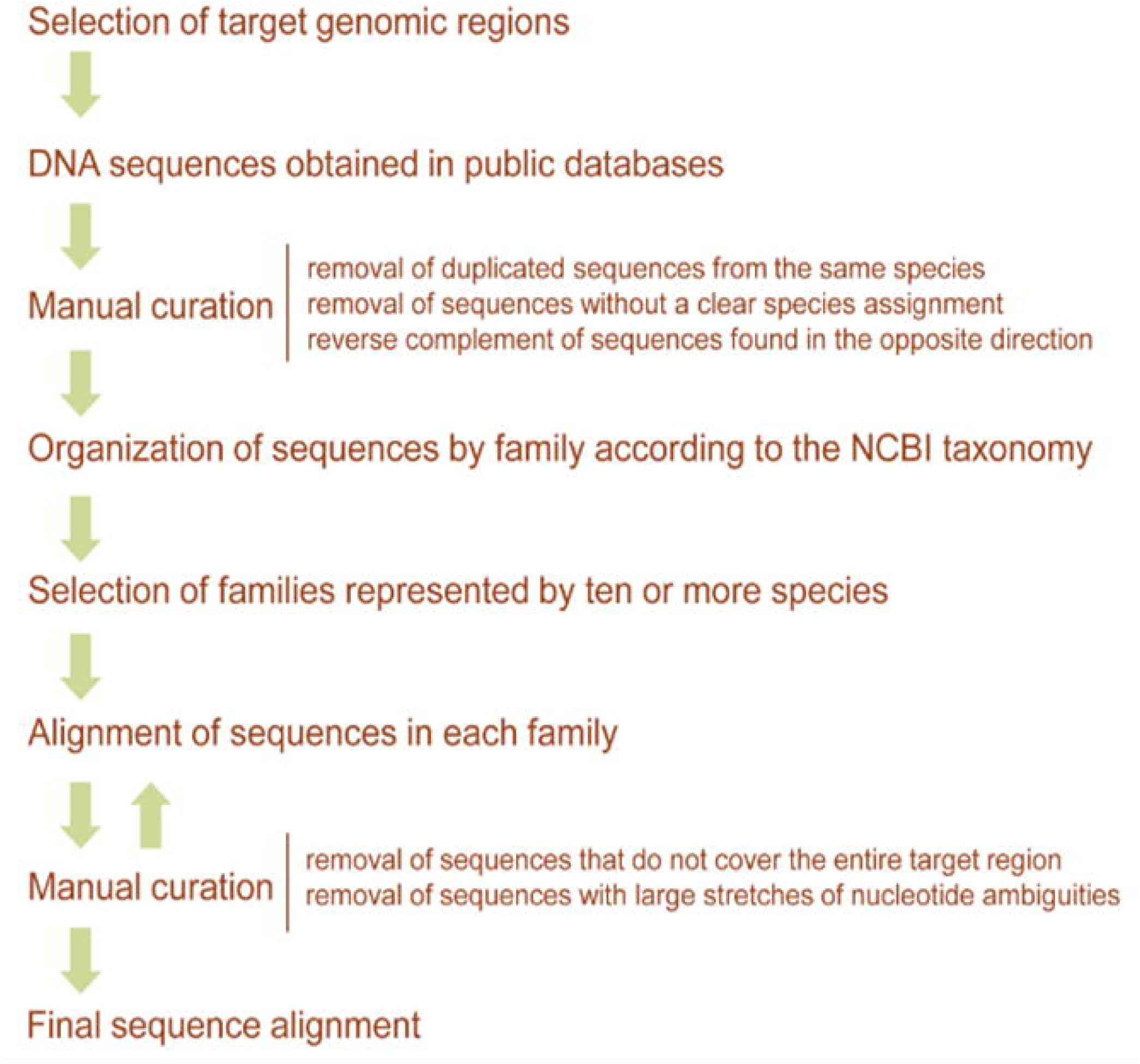
Workflow used to generate the curated alignments, phylogenetic trees, and genetic conservation values stored in PlantAligDB.

## Materials and Methods

### Data curation

We retrieved all nucleotide sequences of different genomic regions from the NCBI Entrez Nucleotide database (http://www.ncbi.nlm.nih.gov) using the Geneious software (Drummond 2009). Different combinations of search terms (e.g. ‘*gene name*’; viridiplantae’; ‘chloroplast’; ‘gene’; ‘complete’) and a maximum limit of 5000 bp as sequence length were used in searches to retrieve the largest number of sequences. The selection was done by eliminating sequences showing one or more of the following features: a) ambiguous name description; b) sequences with high number of nucleotide ambiguities (>50%); c) sequences with large stretches of the region missing (>50%). After a preliminary curation of the data, we selected five cpDNA regions and one nuclear DNA region, which were the most represented in the NCBI database, commonly used in phylogenetic studies for being relevant and informative. The six genomic regions were named according to the gene regions where they are located: *atp*F-*atp*H *(ATPase I subunit – ATPase III subunit), psb*A-*trn*H [*Photosystem II 32 kDa protein – tRNA-His (GUG)*], *trn*L [*tRNA*-Leu (UAA)], *rbc*L (*rubisco large subunit*), *mat*K *(maturase* K) and ITS (internal transcribed spacer). We then removed from the datasets all redundant sequences belonging to the same species and sequences without a clear species assignment. We also reverse complement the sequences that were found in the opposite direction. The sequence orientation for each region is that of the most commonly found in the NCBI database. Therefore, the orientation of the *trn*L *(UAA), atp*F-*atp*H and *rbc*L regions are the same of that used in the reference cpDNA sequence of *Nicotiana tabacum* (NC_001879.2), while the opposite orientation is used for regions *psb*A-*trn*H and *mat*K. The target region named ITS in our database includes the internal transcribed spacer 1, 5.8S rRNA and internal transcribed spacer 2 section of the nuclear ribosomal DNA.

Because a high number of sequences were detected for the *trn*L (UAA) region (more than 50,000 hits), we used the external regions named “C” and “D” and the internal regions named “G” and “H” by (Taberlet *et al*. 2007) as queries in the NCBI Basic Local Alignment Search Tool (BLAST; http://blast.ncbi.nlm.nih.gov/). The search was made against the nucleotide collection (nr/nt) of Tracheophyta (vascular plants) using the Biopython package (www.biopython.org) with an expected threshold of 1000 and a minimum word size of 16. Therefore, our database includes two datasets for the *trn*L (UAA) genomic region: the ‘*trn*L CD’ target region with a length of 577 bp in *N. tabacum*, and the *‘trn*L GH’ with a length of 78 bp in *N. tabacum*, located inside the *trn*L CD region. All information regarding the selected target regions can be found in the *Genomic Regions* section of the PlantAligDB.

The nucleotide sequences of the six regions were organized by family according to the NCBI taxonomy and were aligned (each region and family in separated alignments) using the default parameters of the MUSCLE software (Edgar 2004) running in the Geneious software. The alignment was repeated in some families after excluding sequences that do not cover the entire region of interest and that had large stretches of nucleotide ambiguities. We only used alignments with ten or more species per family to build the PlantAligDB. Some species sequences were lost in this process filter. The neighbor joining phylogenetic tree of each region-family were calculated using Tamura-Nei model using Geneious Tree Builder. The methodology was built in Armadillo Workflow (http://www.bioinfo.uqam.ca/armadillo/) to automate the update process of the database. The latest release update of June 2018 incorporates 514 alignments and phylogenetic trees, from 223 plant families.

#### Conservation measures

The database includes two measures of sequence conservation for each alignment: *percentage of identical sites* (PIS), calculated by dividing the number of identical positions in the alignment for an oligonucleotide by its length and the *percentage of pairwise identity* (PPI), calculated by counting the average number of pairwise matches across the positions of the alignment, divided by the total number of pairwise comparisons.

## Results and Discussion

### Database organization

#### Basic structure

The PlantAligDB database is divided in nine sections (Figure 1): 1) *Home*, provides a brief description of what can be done in the database; 2) *Genomic regions*, describes the regions used in the database; 3) *Taxonomic groups*, the table containing the plant family/region alignments and phylogenetic trees; 4) *BLAST*, search the database using a query sequence by means of the BLASTN algorithm (Altschul *et al*. 1990); 5) *Genetic diversity*, describes the percentage of identical sites (PIS) and pairwise identity values (PPI) for each alignment; 6) *Download*, provides hyperlinks to download the curated alignments; 7) *Tutorials*, contains information about how the database was built, and how to use it; 8) *Citations*; 9) *Contacts*.

#### Taxonomic groups

The database is being regularly updated by our team and currently includes 514 alignments and phylogenetic trees from seven target regions: *atp*F-*atp*H, *psb*A-*trn*H, *trn*L CD, *trn*L GH, *rbc*L, *mat*K and ITS (Table 1). Sequence alignments are provided for 223 different plant families. Currently, the *trn*L GH region has the largest number of sequences (n = 34,674). The *Fabaceae* family has the largest number of aligned species in a target region, with 2599 sequences for the *trn*L GH region. When considering all regions together, the *Fabaceae* (n = 4714), *Poaceae* (n = 4494) and *Asteraceae* (n = 4459) families are those with the highest number of sequences. The alignments for each plant family can be accessed through a dynamic table in the *Taxonomic groups* section of the database by following a hyperlink with the number of species included in each alignment (http://plantaligdb.portugene.com/cgi-bin/PlantAligDB_taxonomicgroups.cgi). The users are able to quickly search and locate a queried feature, order each column using the ascendant or descendent mode, filter the information, download the curated datasets, among other features. The multiple sequence alignment and phylogenetic tree can be visualized by clicking in the number of species present in the alignment.

**Table 1:** Summary of data currently available in the PlantAligDB.

| Target region | Genome | Type | Length (bp) in *Nicotiana tabacum* | Number of alignments | Number of sequences |
| --- | --- | --- | --- | --- | --- |
| *atpF-atpH* | cpDNA | Inter-genic spacer | 502 | 31 | 1025 |
| *psbA-trnH* | cpDNA | Inter-genic spacer | 509 | 79 | 4852 |
| *trnL CD* | cpDNA | Intron | 577 | 44 | 2527 |
| *trnL GH* | cpDNA | Intron | 78 | 173 | 34674 |
| *rbcL* | cpDNA | Protein-coding gene | 1434 | 39 | 1748 |
| *matK* | cpDNA | Protein-coding gene | 1530 | 113 | 11341 |
| **ITS** | nuDNA | Transcribed spacers and 5.8S gene | 678 | 35 | 9885 |
| *Total* |  |  |  | 514 | 66052 |

#### Genetic diversity

The PIS values in our current dataset vary from 0.16% to 99.07% (Table 2). Nevertheless, the PIS and PPI sequence conservation measures are not intended to be used for comparison of different families and/or regions, since the number of sequences in each alignment can be very different. The results must be interpreted at the light of the number of sequences and the representativeness of the sequences included in the alignment file. The *mat*K was the region with the lowest PIS value (0.16%) [Figure 2. f)], while the *trn*L GH was the region with the highest PIS value (99.07%), as can be seen in Figure 2 d) and Table 2. The *rbc*L was the most conserved region [Figure 2 e)] with an average of 78.48%, while the ITS was less conserved with an average of 28.84% (Table 2). Our results are in accordance with earlier studies where *atp*F*-atp*H and *psbA-trn*H were found to be more variables than *mat*K (Lahaye 2008). The *trnL* CD regions showed values slightly more conserved than *atp*F*-atp*H [Figure 2 c) and a)]. The lowest PPI value (69.96%) was found in *psb*A*-trn*H region [Figure 2 b)] and the highest was 100% in *trn*L GH, as shown in Figure 2 d) and Table 2. The ITS region was less conserved with an average of 87.21% [Table 3 and Figure 2 g)], while the *trn*L GH was the most conserved with an average of 97.09% [Table 3 d)].

**Figure 2:**
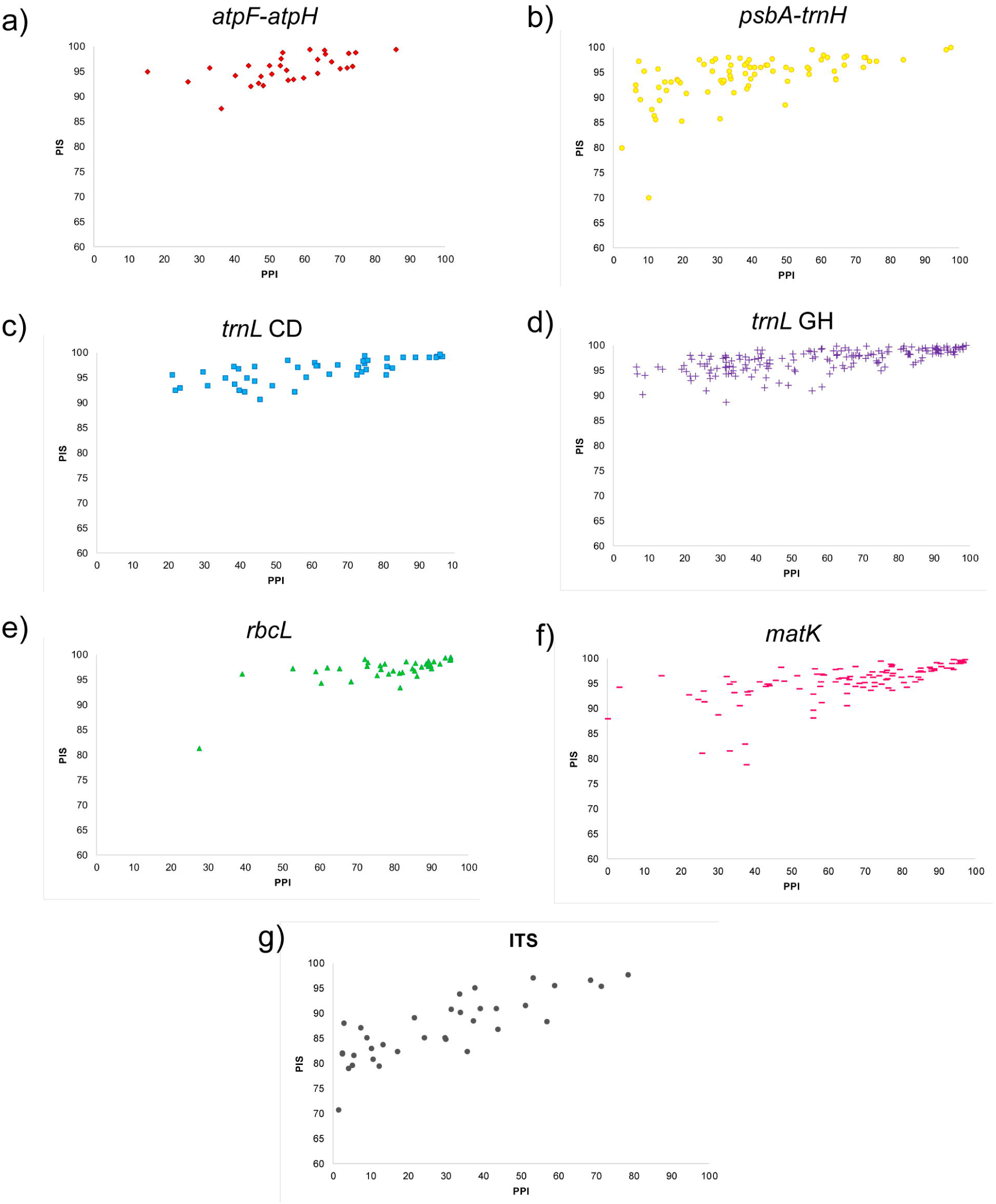
Graphic representation of the measures of sequence conservation PPI and PIS for each region-family alignment: a) *atp*F*-atp*H region b), *psb*A*-trn*H region, c) *trn*L CD region, d) *trn*L GH region, e) *rbc*L region, f) *mat*K region and g) ITS region.

**Table 2:** Average percentage of identical sites (PIS) values in all plant families organized by genomic region.

| PIS | *atpF-atpH* | *psbA-trnH* | *trnL CD* | *trnL GH* | *rbcL* | *matK* | **ITS** |
| --- | --- | --- | --- | --- | --- | --- | --- |
| **Mean** | 55.05 | 39.1 | 61.13 | 58.62 | 78.48 | 65 | 28.84 |
| **Max** | 85.95 | 97.42 | 96.46 | 99.07 | 95.24 | 97.3 | 78.45 |
| **Min** | 15.19 | 2.3 | 21 | 6.43 | 27.47 | 0.16 | 1.13 |

**Table 3:**
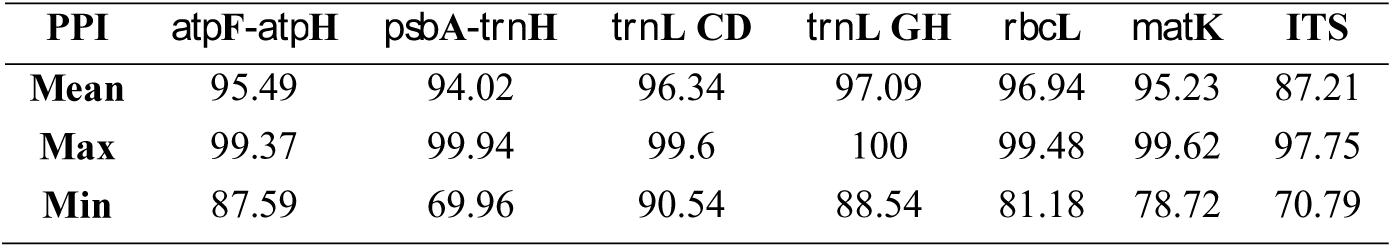
Average percentage of pairwise identity (PPI) values in all plant families organized by genomic region.

| PPI | *atpF-atpH* | *psbA-trnH* | *trnL CD* | *trnL GH* | *rbcL* | *matK* | **ITS** |
| --- | --- | --- | --- | --- | --- | --- | --- |
| **Mean** | 95.49 | 94.02 | 96.34 | 97.09 | 96.94 | 95.23 | 87.21 |
| **Max** | 99.37 | 99.94 | 99.6 | 100 | 99.48 | 99.62 | 97.75 |
| **Min** | 87.59 | 69.96 | 90.54 | 88.54 | 81.18 | 78.72 | 70.79 |

#### Sequence alignments and phylogenetic trees

The alignments stored in the database can be visualized using a dynamic browser-based application named Wasabi (http://wasabiapp.org/) (Veidenberg *et al*. 2016) with multiple options for the visualization and analysis of sequence data and phylogenetic trees. This resource is particularly useful to help researchers selecting the most appropriated genomic regions for their investigations. The users can zoom in and out the selected regions of the alignments, collapse regions with gaps, alternate between column and row selection, remove or add sequences, realign sequences, and export the sequence data in the FASTA format. If an Wasabi account is created, the user can re-align specific alignments with PAGAN (Loytynoja *et al*. 2012) and PRANK (Loytynoja 2014). The user can merge different alignments from identical regions (Hollingsworth et al. 2009) and different families using the PAGAN application. The download of the complete database of curated alignments is accessible in the *Download* section of the database.

#### How to use the PlantAligDB

To explore a genomic region, a researcher must start by accessing the ‘Genomic Regions’ section in the menu bar. For example, by selecting *psb*A*-trn*H, the user will find a brief description of the genomic region and the name, size and position of that region in the reference genome, and the families with available alignments. The user should select the ‘Taxonomic Groups’ section in the menu bar to search for a specific plant family. Then, through the search tool on the right side of the page, the user can type the name of the family and a dynamic table will be displayed with the number of sequences for each region. The user can also access a hyperlink with the description of the family and taxonomic tree using the resource Tree of Life Web Project (http://tolweb.org). Clicking on the number of sequences, the database is redirected to the Wasabi tool. The user can create a Wasabi account by providing an e-mail or choosing a temporary account, which allows to realign sequences with PAGAN or PRANK. The user can merge a PAGAN realignment with alignments of other families on the same region, by selecting a file on his local computer in the “alignment extension” option.

#### Availability and design

The PlantAligDB is freely available at http://plantaligdb.portugene.com and is optimized for the major web browsers (Internet Explorer, Firefox, Safari, and Chrome). The SQLite local database is used for data storage and runs on an Apache web server. The dynamic HTML pages were implemented using CGI-Perl and JavaScript and the dataset table views were generated using the JQuery plugin DataTables v1.9.4 (http://datatables.net/). The PlantAligDB visualization tables are generated automatically. The process of database update is optimized for large datasets. There are no access restrictions for academic and commercial use.

## Acknowledgments

This work was supported by the Portuguese Foundation for Science and Technology (FCT), European Regional Development Fund (ERDF) and Operational Human Potential Program [IF/01356/2012 to F.P. and SFRH/BPD/100912/2014 to J.C.]. J.C. also acknowledges the FCT funding for his research contract at CIIMAR, established under the transitional rule of Decree Law 57/2016, amended by Law 57/2017. C.S. was supported by the “Programa Ciências Sem Fronteiras” from the Conselho Nacional de Desenvolvimento Científico e Tecnológico (CNPq) and Coordenação de Aperfeiçoamento de Pessoal de Nível Superior (CAPES), Brazil. CIIMAR was partially supported by the Strategic Funding UID/Multi/04423/2019 through national funds provided by FCT and ERDF in the framework of the programme PT2020.

## Author Contributions

CS, JC and FP contributed equally to designed and performed research, analysed data and wrote the paper.

